# Antero-posterior patterning in the brittle star *Amphipholis squamata* and the evolution of body plans across echinoderms

**DOI:** 10.1101/2025.02.13.638179

**Authors:** L. Formery, P. Peluso, D. R. Rank, D. S. Rokhsar, C. J. Lowe

**Affiliations:** Department of Biology, Hopkins Marine Station, Stanford University, Pacific Grove, CA, USA; Department of Cell and Molecular Biology, University of California Berkeley, Berkeley, CA, USA; Pacific Biosciences, Menlo Park, CA, USA; Molecular Genetics Unit, Okinawa Institute of Science and Technology, Onna, Okinawa, Japan; Chan Zuckerberg BioHub, San Francisco, CA, USA

**Keywords:** Echinoderms, Axial patterning, Body plan evolution, Pentaradial symmetry, Hox genes

## Abstract

Although the adult pentaradial body plan of echinoderms evolved from a bilateral ancestor, identifying axial homologies between the morphologically divergent echinoderms and their bilaterian relatives has been an enduring problem in zoology. The expression of conserved bilaterian patterning genes in echinoderms provides a molecular framework for resolving this puzzle. Recent studies in juvenile asteroids suggest that the bilaterian antero-posterior axis maps onto the medio-lateral axis that is perpendicular to each of the five rays of the pentaradial body plan. Here we test this hypothesis in another echinoderm class, the ophiuroids, using the cosmopolitan brittle star *Amphipholis squamata*. Our results show that the general principles of axial patterning are similar to those described in asteroids, and comparisons with existing molecular data from other echinoderm taxa support the idea that medio-lateral deployment of the AP patterning program across the rays predates the evolution of the asterozoan and likely the echinoderm crown-groups. Our data also reveal expression differences between *A. squamata* and asteroids, which we attribute to secondary modifications specific to ophiuroids. Together, this work provides important comparative data to reconstruct the evolution of axial properties in echinoderm body plans.

## Introduction

Echinoderms are a phylum of marine invertebrates comprising five extant classes: crinoids (sea lilies), holothuroids (sea cucumbers), echinoids (sea urchins and sand dollars), asteroids (sea stars) and ophiuroids (brittle stars and basket stars). Adults of all classes are characterized by the presence of a calcitic endoskeleton, a water vascular system, and pentaradial symmetry [1]. Molecular phylogenies consistently support echinoderms as being sister-group to hemichordates [2–4], indicating that the pentaradial organization of their adult body plan evolved through axial reorganization of a bilateral ancestor. Despite the rich fossil record of echinoderms, the nature of this axial reorganization remains enigmatic, and morphological comparisons between pentaradial echinoderms and their bilateral relatives have been historically challenging [1,5]. Recently, the investigation of conserved molecular developmental programs involved in bilaterian axial patterning have begun to provide some insights into the evolutionary origins of the echinoderm pentaradial body plan [6,7]. Among these programs is the antero-posterior (AP) patterning program, a conserved suite of genes that patterns the AP axis of the ectoderm in animal groups as distantly related and morphologically divergent as arthropods, annelids, hemichordates or chordates [8–13]. AP patterning genes include transcription factors involved in the patterning of anterior (head) territories, along with the Hox complex that is deployed in posterior territories and controls trunk patterning. In bilaterians, this suite of transcription factors is regulated by a Wnt gradient set up by the interaction of posteriorly expressed ligands and their antagonists localized anteriorly [14–18]. The exquisite conservation of this patterning program across diverse bilaterian body plans offers a robust molecular readout of the AP axis [19], which has potential to unravel cryptic axial properties that might have been masked by the echinoderm divergent morphology.

Several models have been proposed for the deployment of the AP patterning program in the pentaradial body plan of echinoderms, based on a combination of morphological, paleontological and molecular data. The duplication hypothesis postulates that each of the five echinoderm rays arise from consecutive duplications of the ancestral AP axis [20,21], and implies staggered expression of AP markers along the proximo-distal axis of the rays. Alternatively, the stacking model proposes that the ancestral AP axis is homologous to the oral-aboral axis of adult echinoderms [5,22,23]. This model is based on the reorganization of the larval coelomic compartments during the formation of the adult body plan, and on sequential Hox gene expression across the coelomic compartments stacked along the oral-aboral axis of the animal in several echinoderm species [24–27].

More recently, a comprehensive survey of AP patterning markers in *P. miniata* led to the proposal of the new “ambulacral-anterior” model [6]. In this model, the midline of the ambulacral ectoderm, which consists mostly of the radial nerve cords along each ray, displays the molecular identity of the most anterior bilaterian regions expressing genes such as *hedgehog*, *sfrp1/5*, *fzd5/8*, *six3/6* and *nkx2.1* (hereafter referred to as anterior head markers). These markers are expressed in the proboscis and forebrain of hemichordates and vertebrates, respectively. Ambulacral regions, located on the lateral sides of the midline and comprising the ectoderm wrapping around the tube feet, express genes with the most caudal limit of expression in the posterior head ectoderm in other deuterostomes, such as *irx*, *dmbx*, *otx*, *barH* and *pax6* (hereafter referred to as posterior head markers). In hemichordates and vertebrates this territory corresponds to the collar and midbrain, respectively. Finally, expression of genes that mark the boundary between the head and trunk in bilaterians such as *gbx, pax2/5/8* and *hox1* (hereafter referred to as head-trunk boundary markers) is detected at the margin of the ambulacral ectoderm abutting the interradial epidermis. Surprisingly, an ectodermal territory corresponding to the bilaterian trunk defined by the expression of the remaining Hox genes is absent, and suggests that from an ectoderm patterning perspective, asteroids are essentially head-like animals. Posterior Hox genes are still expressed in internal germ layers, but are uncoupled from the axial polarity of the ectoderm and follow an independent patterning logic [6,7,28]. The medio-lateral deployment of anterior patterning genes across the ambulacral ectoderm of *P. miniata* is largely congruent with molecular data from the echinoid *P. japonica* [7], but still has to be formally tested in other echinoderm classes.

Ophiuroids (brittle stars), examined below, are the sister taxon of asteroids [29,30]. While ophiuroids superficially resemble asteroids owing to their stellate body plan, they exhibit substantial differences at the anatomical and developmental levels. Unlike asteroids, they have a blind gut lacking an anus; a madreporite located orally and embedded in the mouth skeleton; a flattened central disc clearly offset from the arms; and the arms themselves are highly articulated and flexible [31]. Although the rays of all echinoderm classes are metameric [32], this character is particularly pronounced in ophiuroids, whose arms are constructed as a succession of discrete brachial segments. The ambulacra in ophiuroids and both the circumoral nerve ring and the radial nerve cords are subepidermal, in contrast with asteroids where the radial nerve cords are embedded within the epidermis at the bottom of ambulacral grooves [33–35]. Most of our knowledge on gene expression in the pentaradial body plan of ophiuroids come from studies of *Amphiura filiformis* [36–38], but there is no comprehensive study of axial patterning genes during adult body plan development.

Here, we investigate the deployment of the bilaterian AP patterning system in juvenile stages of the brittle star *Amphipholis squamata* (Delle Chiaje, 1828) (Figure 1A). *A. squamata* is a cosmopolitan species complex of small ophiuroids characterized by simultaneous hermaphroditism, a brooding life style, and population-specific genome duplication and/or allopolyploidy events [39–41]. We start by describing the morphological development and juvenile anatomy of *A. squamata*, before comprehensively surveying the expression of 21 AP patterning genes within the juvenile body plan. We find strong parallels between AP patterning gene expression in *A. squamata* and *P. miniata* that support extending the ambulacral-anterior model to ophiuroids. We also report gene expression differences between *A. squamata* and other echinoderm species, which we attribute to secondary modifications arising in the ophiuroid stem lineage. Our findings help differentiate between developmental innovations along the echinoderm stem involved in the establishment of the radial body plan, and those involved in the later diversification of echinoderm crown-groups into more specialized forms.

**Figure 1:**
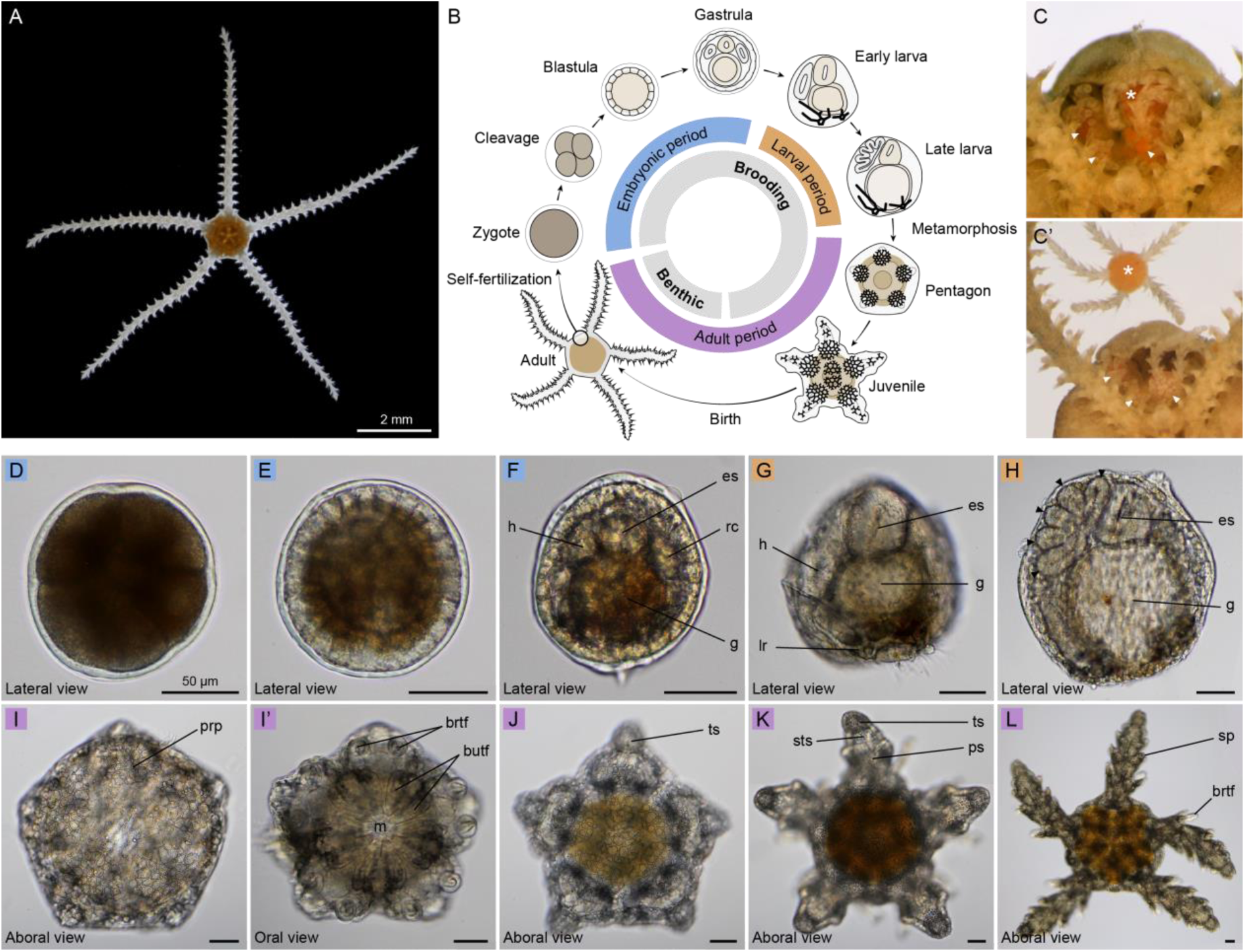
Morphological development of *Amphipholis squamata*. **A**, Adult *Amphipholis squamata*, viewed from the oral side. **B**, Schematic representation of the life cycle in *A. squamata*. **C**, **C’**, Images of a dissected adult showing four distinct juveniles (arrowheads and asterisk) developing within the bursa (**C**), with the largest (asterisk) being manually extracted (**C’**). **D**-**L**, selected developmental stages of *A. squamata*: cleavage (**D**), blastula (**E**), gastrula (**F**), early larva (**G**), late larva (**H**), pentagon (**I**), early juvenile (**J**), mid-juvenile (**K**), late juvenile (**L**). The pentagon stage is shown in aboral (**I**) and oral (**I’**) view, other juvenile stages are shown in aboral views (**J**-**L**). butf: buccal tube foot, brtf: brachial tube foot, es: esophagus, g: gut, h: hydrocoel, lr: larval skeletal rod, prp: primary radial plate, ps: proximal segment, rc: right coelom, sp: spine, sts: sub-terminal segment, ts: terminal segment. Scale bars: 2 mm (**A**), 50 µm (**D**-**L**).

## Results

### Morphological development of *Amphipholis squamata*

Fertilization in *A. squamata* occurs internally within the bursal sacs of the adult [42], and likely involves high rates of selfing [43]. Subsequent development also takes place internally, making it difficult to observe directly. Embryos develop inside the adult bursal sacs up until late juvenile stages, at which point they crawl out of the bursae through the bursal slits. Morphogenesis has been described in detail by Fell [44], but we started here by re-investigating this process using modern microscopy. As in other indirect-developing echinoderms, the development in *A. squamata* can be divided into three periods: embryonic, larval and adult (Fig.1B). To characterize developmental stages, we dissected adult bursal sacs and extracted asynchronously developing individuals (Fig. 1C,C’; Supplementary material 1). Since there were no external indications of the number or developmental stages of the progeny developing within the bursae, the likelihood of finding a particular developmental stage was proportional to the duration of that stage. Embryonic development is likely rapid, since these stages were rarely encountered; the more commonly observed juvenile stages apparently lasted several days or weeks. When dissecting a sample of twenty adults, we collected 217 developing progeny, including 8.7% in embryonic stages, 8.2% in larval stages, and 82.9% in juvenile stages. While embryos and juveniles developed freely within the bursae, larvae were embedded into the bursal walls and had to be torn apart from adult tissues. Owing to their small adult size, minimal husbandry requirements, and profusion of juvenile stages developing within each adult, *A. squamata* was thus an ideal species to investigate post-metamorphic development in ophiuroids.

The embryonic period in *A. squamata* included typical cleavage, blastula and gastrula stages (Fig.1D-F). All the embryonic stages were densely pigmented (Fig.1D), with blastomeres uniformly pigmented during the zygote and cleavage stages and pigments becoming progressively restricted to the basal side of the blastomeres in the blastula stage (Fig.1E). At the gastrula stage, pigments become restricted to the posterior endoderm (Fig. 1F). Cell division patterns during cleavage and germ layers formation during gastrulation could not be observed directly given the scarcity of embryonic stages found by dissecting adults. However, gastrulation resulted in the typical segregation of the embryo into an outer ectoderm layer, an inner endoderm layer divided into esophagus and stomach, and a mesoderm layer comprising a left and right coelomic pouch, the left one being identified as the hydrocoel (Fig.1F). Following embryogenesis, the larval morphology appeared greatly modified compared to pelagic ophiopluteus larvae of other ophiuroids, presumably as a consequence of the highly derived viviparous brooding life history in *A. squamata*. Yet, the larval period was marked by the presence of an esophagus that underwent muscular contractions, reminiscent of the functional digestive tract of typical ophioplutei (Supplementary material 2). Larvae were initially bilaterally symmetrical, but the hydrocoel on the left side quickly grew in size while the right coelom was reduced, resulting in marked left-right asymmetry (Fig. 1G). During the larval period two vestigial skeletal rods developed at the posterior end of the larva, but never extended into distinguishable arms as in pelagic ophioplutei. At the late larval stage, five lobes eventually budded out of the hydrocoel, constituting the anlage of the future radial canals of the adult (Fig.1H). Metamorphosis was not observed and likely happened rapidly. Following metamorphosis, individuals exhibited the definitive pentaradial symmetry, and had a simple pentagon shape lacking any visible arms (Fig.1I). Characteristic fenestrated skeletal plates appeared on the aboral side of the animal, starting with the five primary radial plates (Fig.1I). The mouth opened at the center of the oral surface and was surrounded by five pairs of buccal tube feet (Fig.1I’). At the same time, the first pairs of brachial tube feet formed on both sides of the pentagon radii (Fig.1I’). Arm elongation progressively became evident as the vertices of the pentagon started to protrude outward with the formation of the terminal arm segment, which eventually separated the disk from the growing arms (Fig.1J). After the formation of the terminal segments, additional segments were intercalated sub-terminally within each arm (Fig.1J-L). Besides the 5 terminal segments, all newly intercalated segments were morphologically identical, and kept growing over time, so that the largest segments were located closest to the disk. Each segment supported a pair of brachial tube feet located at the distal joint, and up to three pairs of spines that became evident in oldest segments. According to the definition provided by Hyman [1], these tube feet define the position of the ambulacra along the midline of each ray in ophiuroids. No juveniles with more than seven arm segments were found, indicating that they exited the adult bursae around this stage.

### *Amphipholis squamata* juvenile anatomy

To provide an anatomical reference for describing the expression of AP patterning genes in *A. squamata* juveniles, we first used a combination of chemical staining, immunostaining, and fluorescent *in situ* hybridization to highlight the organization of the juvenile endoskeleton, muscles, water vascular system and nervous system. To describe these structures we followed established terminology for *A. squamata* [44–46] and other ophiuroids [1,47]. We surveyed three distinct stages of juvenile development that we identified by the number of arm segments and referred to as early (0-1 segment), mid-(2-3 segments) and late (>3 segments) juveniles (Fig.1J-L). To obtain fluorescent *in situ* hybridization chain reaction (HCR) probes for anatomical and AP patterning markers, we generated long-read (PacBio Iso-Seq) RNA sequencing from adult tissues. Although we did not determine the number of genome copies present in the population of *A. squamata* used for this study, we reasoned that multiple copies of the genes investigated by HCRs could be present. However, since different populations of *A. squamata* share almost identical sequence variants with only little subsequent divergence [41], we hypothesized that our HCR probes would cross react with all recent paralogues, showing composite expression from all sequence variants. To verify this assumption as a preliminary step for other HCR assays, we designed distinct probe sets against four sequence variants of the transcription factor *nkx2.1* and observed identical expression patterns (Supplementary material 3; Supplementary material 4), corroborating that functional divergence across recent paralogues are unlikely at the time scale of *A. squamata* polyploidization events.

To describe skeletal anatomy, we labelled the endoskeleton in *A. squamata* juveniles by incubating animals in calcein, a fluorescent calcium analogue that is incorporated into the skeletal matrix during growth [48]. In early juveniles, the teeth were visible on the oral side of the disk, in addition to a series of plates associated with the jaw apparatus (Fig.2Aa). The general organization of the skeletal plates on the oral side of the disk remained similar at later stages, although each plate grew considerably (Fig.2Aa). On the aboral side, the disc was completely covered by a series of circular concentric plates (Supplementary material 5). The arm skeleton was organized into repeated sets of ossicles within each segment, comprising four shield plates and a pair of vertebrae (Fig.2Ab-Ad). Shield plates developed just beneath the epidermis and included two lateral shields, which at late stages supported the spines, one oral shield, and one aboral shield. At the early juvenile stage, the first set of oral shield plates was located at the base of each prospective arm, between the first set of brachial tube feet, while developing lateral shield plates were visible on either side of the first arm segments (Fig.2Aa). By the late juvenile stage, the shield plates completely enclosed the tissues of the oldest segments, with the exception of the protruding brachial tube feet. By contrast, the vertebrae were internal ossicles that developed deeper within the aboral half of the arm tissues (Fig.2Ab-Ad). They had a thin, elongated shape and connected the proximal and distal end of each segment where they were slightly enlarged.

**Figure 2:**
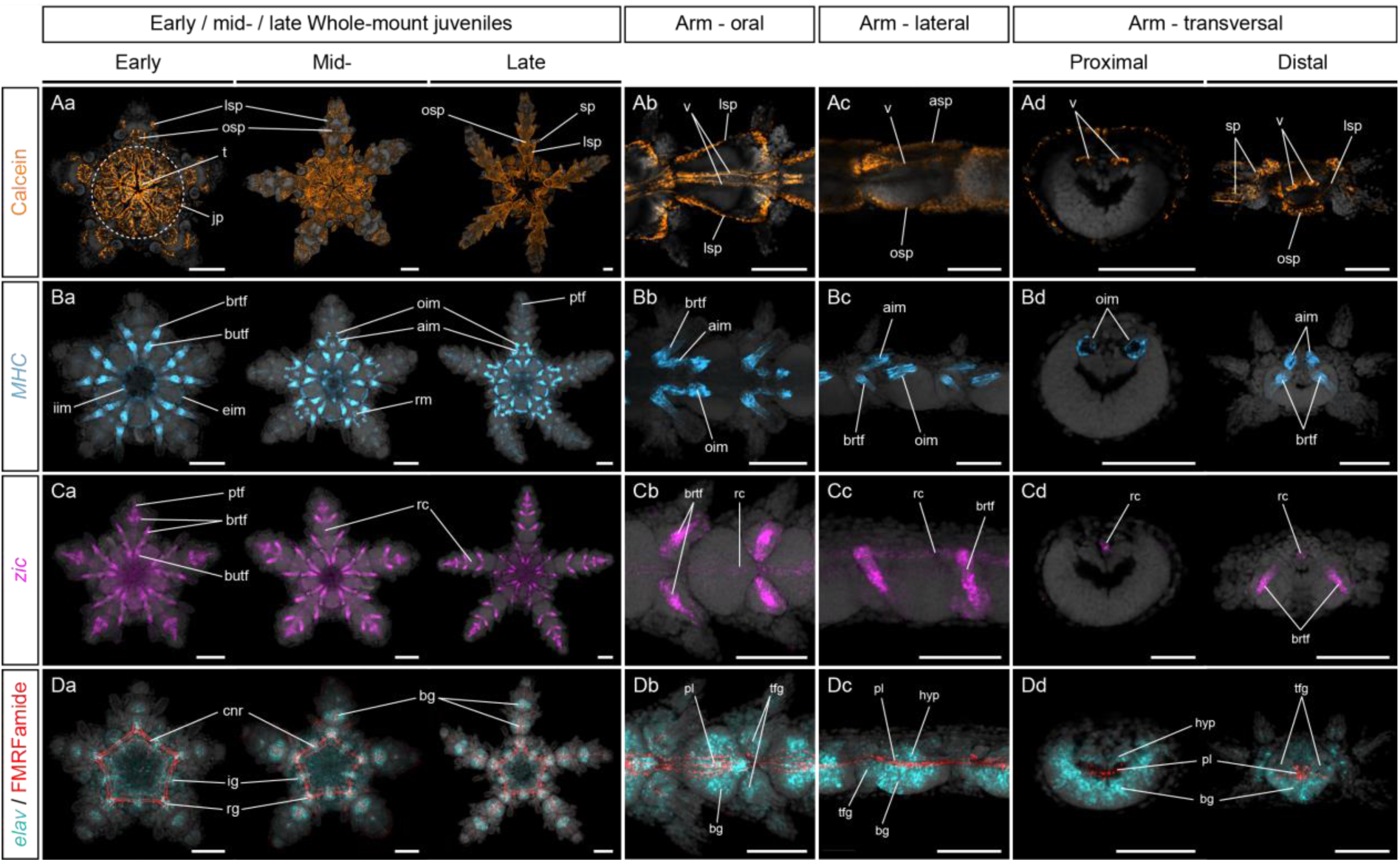
Anatomy of *Amphipholis squamata* juveniles. Calcein stainings (**A**), single HCRs for *MHC* (**B**), *zic* (**C**) and HCRs for *elav* combined with immunostainings against FMRF-amide-like neuropeptide (**D**) showing the anatomy of the endoskeleton (**A**), muscles (**B**), water vascular system (**C**) and nervous system (**D**) in *Amphipholis squamata* whole-mount early (left panel), mid-(middle panel) and late (right panel) juveniles viewed from the oral side (**Aa**, **Ba**, **Ca**, **Da**), and in detailed oral (**Ab**, **Bb**, **Cb**, **Db**), lateral (**Ac**, **Bc**, **Cc**, **Dc**) and transversal (**Ad**, **Bd**, **Cd**, **Dd**) views of brachial segments. Transversal views show section through the proximal (left panel) and distal (right panel) regions of a brachial segment. All samples are counterstained with DAPI (grey) to mark cell nuclei. aim: aboral intervertebral muscle, asp: aboral shield plate, bg: brachial ganglion, brtf: brachial tube foot, butf: buccal tube foot, cnr: circumoral nerve ring, eim: external interradial muscle, hyp: hyponeural neuroepithelium, ig: interradial ganglion, iim; internal interradial muscle, jp: jaw plates, lsp: lateral shield plate, oim: oral intervertebral muscle, osp: oral shield plate, pl: plexus, ptf: primary tube foot, rc: radial canal, rg: radial ganglion, rm: radial muscle, sp: spine, t: tooth, tfg; tube foot ganglion, v: vertebra. Scale bars: 100 µm.

To examine the muscular anatomy of *A. squamata* juveniles we used fluorescent *in situ* hybridization chain reactions (HCR) with probes corresponding to *Myosin Heavy Chain* (*MHC*) (Supplementary material 6), which has been used as a muscle marker in several echinoderms [49,50]. In all three juvenile stages, HCRs revealed strong *MHC* expression in the mesoderm lining of the tube feet, indicating the muscular nature of this epithelium, as reported for other echinoderms [51,52] (Fig.2Ba). *MHC* expression in the tube feet was most obvious in the buccal tube feet and the first set of brachial tube feet, but only became apparent at later stages in the primary tube feet that constituted the terminal end of the water vascular system within each arm. At the early juvenile stages, the other tissues expressing *MHC* were restricted to the disk and corresponded to the external and internal interradial muscles that are parts of the jaw apparatus, and to the epithelial lining of the esophagus (Fig.2Ba). At the mid-juvenile stage, *MHC* started to be expressed in the radial muscles located at the base of each arm (Fig.2Ba). In addition, two pairs of intervertebral muscles connecting each arm segment become visible as arms elongate in the mid- and later juvenile stages (Fig.2Ba-Bd). These include the aboral intervertebral muscles, which are located at the distal end of each segment, and the oral intervertebral muscles, which are located in the proximal region of the next segment (Fig.2Ba-d). Both intervertebral muscles appear as longitudinal bundles of *MHC*+ fibers encircling the region occupied by the extremities of the vertebrae (Fig.2Ba-d). No muscular structure was present on the aboral side of the disk.

To investigate the structure and distribution of the water vascular system, we investigated the expression of the transcription factor *zic* (Supplementary material 6), which is expressed in the hydrocoel in echinoids [7]. As expected, *zic* expression in *A. squamata* juveniles was restricted to the water vascular system and was consistent through all three stages investigated (Fig.2Ca). *Zic* was highly expressed in the mesoderm lining of the tube feet, similar to *MHC*, although unlike *MHC* its expression in the primary tube feet was already clearly visible at the early juvenile stage. *Zic* was also expressed in the radial canals running along the midline of each arm on the aboral side, between the spaces occupied by the two vertebrae (Fig.2Ca-d). Expression was higher in the distal half of the radial canal than in the proximal part. Of note, *zic* was not expressed in the ring canal.

Finally, to visualize the nervous system, we used HCRs to survey the expression of the neuronal marker *elav* to label cell bodies [53], in combination with an antibody targeting FMRFamide-like neuropeptides to label neurites [54]. The nervous system in adult echinoderms consists of five radial nerve cords running along the five rays linked by a circumoral nerve ring that encircles the pharynx of the animal [1,34,47,52]. In addition, there are peripheral structures such as the lateral nerves innervating the tube feet, and a plexus below the epidermis of the animal. The radial nerve cords and the circumoral nerve ring, which are the most prominent neural structures, have historically been defined by a large ectoneural component on the oral side and a thinner hyponeural layer on the aboral side, mostly associated with motor functions [55]. At the early juvenile stage, *elav* was broadly expressed in the oral epidermis of the disk (Fig.2Da). Concentrations of *elav*+ nuclei marked five interradial ganglia located between arms, and five radial ganglia located at the base of each arm (Fig.2Da). Together, these ten ganglia were connected by a plexus of FMRFamide-like positive neurites forming the circumoral nerve ring (Fig.2Da). At this stage, the circumoral nerve ring was pentagonal in shape and was formed by several concentric tracts of neurites. In each radius, FMRF-amide-like-positive neurites branched out from the circumoral nerve ring to form the most proximal part of the plexus of the radial nerve cords, but did not extend yet to the tip of the developing arms. The radial nerve cords became evident at later stages, as the arms elongated, and were characterized by a prominent neuroepithelium running along the arm midlines down to the terminal segment, which we identified as the ectoneural subsystem of the radial nerve cords. In the ectoneural subsystem of *A. squamata*, *elav*+ nuclei defined large ganglion-like swellings of the neuroepithelium in the proximal/oral half of each arm segment, which we referred to as the brachial ganglia (Fig.2Da-d). In transverse views, these ganglia exhibited a crescent-shape lining the oral epidermis of the arms, and were made of multiple layers of densely packed cell bodies (Fig.2Dd). As shown by nuclear staining, the ectoneural neuroepithelium in the distal part of each segment narrowed and connected to the brachial ganglia of the next segment, although cell bodies in the interganglionic region did not express *elav* (Fig.2Db,c). In addition to the ectoneural neuroepithelium, several parallel tracts of FMRF-amide-like positive neurites formed a plexus occupying a crescent-shaped groove located roughly halfway between the oral and aboral surface of the arms and overlying the ectoneural neuroepithelium (Fig.2Da-d). *Elav*+ nuclei evidenced another, smaller neuroepithelium located directly above the plexus and below the radial canals and intervertebral muscles, which we identified as the hyponeural subsystem of the radial nerve cords. Together, the ectoneural neuroepithelium, the plexus and the hyponeural neuroepithelium constituted the radial nerve cords within each arm. At these juvenile stages, the radial nerve cords represented by far the largest anatomical structure, while in adult specimens the muscles typically occupy a much larger relative space [45]. In addition to the radial nerve cords, other neural structures were observed. These included scattered *elav*+ neurons in the epidermis of the disk, tube feet and spines. Finally, lateral projections of the radial nerve cords in the distal part of each segment encircled the stem of each brachial tube foot, forming podial ganglia (Fig.2Da-d).

With this anatomical understanding in hand, we next investigated the expression pattern of AP patterning genes in *A. squamata* juveniles. A recent survey in the asteroid *P. miniata* examined the expression of 36 AP patterning genes and established that AP patterning genes in *P. miniata* fall into four distinct categories based on their region of expression: (1) anterior head markers predominantly expressed along the midline of the ambulacral ectoderm, (2) posterior head markers predominantly expressed in the epidermis covering the tube feet, (3) head-trunk boundary markers predominantly expressed at the boundary between the ambulacral and interradial ectoderm, and (4) genes expressed in internal germ layers (endoderm and mesoderm) [6]. Of these, we shortlisted the most relevant for comparative purposes based on their relative overlapping expression patterns. We then retrieved the cDNA sequences of 21 of these genes from the Iso-Seq dataset (Supplementary material 6) and synthesized HCR probes to survey their expression in *A. squamata*. In the next sections, we report the expression domains of these genes in *A. squamata* and compare with published data from other echinoderm species.

### Expression of anterior head markers

Anterior head markers are predominantly expressed in the circumoral nerve ring and the radial nerve cords in *P. miniata*, even though their individual expression domains within these territories show marked differences. These genes include the ligand *hedgehog*, which is involved in patterning the telencephalon in vertebrates [56,57] and the tip of the proboscis ectoderm in hemichordates [12,58], as well as components of the Wnt pathway such as *fzd5/8* and *sfrp1/5* that in other bilaterian have conserved roles in patterning the most anterior territories [15,17,59]. This category also includes several transcription factors such as *nkx2.1* and *six3/6*, which in vertebrates and hemichordates are expressed during the development of the forebrain and the proboscis, respectively [12,57,60,61]. We started to analyze AP patterning in *A. squamata* by determining the expression patterns of these five medial ambulacral ectoderm genes using HCRs in early, mid-and late juveniles.

First, we found that *hedgehog* was detected at low levels at the early juvenile stage in a spot of the oral epidermis located at the base of each developing arm (Fig.3Aa). Later, at the mid- and late juvenile stages, *hedgehog* expression extended in a narrow territory along the oral midline of the ectoneural part of the radial nerve cords. Additional expression domains were detected branching laterally from the midline at the base of each pair of brachial tube feet (Fig.3Aa,Ab). *Hedgehog* expression was restricted to the oral layers of cells within the ectoneural neuroepithelium, and did not reach the aboral layers in contact with the plexus (Fig.3Ac,Ad). Overall, *hedgehog* expression pattern in *A. squamata* was strikingly similar to that described in *P. miniata*, where it is also expressed along the midline of the radial nerve cords and the base of each tube foot [6].

We next surveyed the expression of an antagonist (*sfrp1/5*) and receptor (*fzd5/8*) of the Wnt signaling pathway. In *A. squamata sfrp1/5* was consistently expressed at the early, mid- and late juveniles in the oral region of the disk, and in a thin stripe orally along the midline of the ectoneural part of the radial nerve cords (Fig.3Ba,Bb). Like *hedgehog*, *sfrp1/5* showed the most medial expression pattern of all the genes investigated in this study, and perfectly delineated the midline of the arms. The *sfrp1/5* expression domain was thicker on the oral side than the aboral side, but unlike *hedgehog* spanned the entire oral-aboral extent of the ectoneural ganglia and connected with the overlying plexus (Fig.3Bb,Bc). In addition to the expression domain along the midline of the radial nerve cords, a group of cells expressing *sfrp1/5* was detected within each arm segment on either side of the midline in the distal part of the brachial ganglia (Fig.3Bb), and also outside of ectodermal derivatives in the radial canal, which is also located along the arm midline (Fig.3Bd). Like *sfrp1/5, fzd5/8* was also predominantly expressed in the medial region of the ectoneural part of the radial nerve cords and in the circumoral nerve ring at the three stages examined (Fig.3Ca). Unlike *sfrp1/5*, however, its expression was exclusively restricted to the oral layers of the ectoneural neuroepithelium but extended more laterally than that of *hedgehog* or *sfrp1/5* (Fig.3Cb-Cd). *Fzd5/8* could also be detected outside the radial nerve cords in mesoderm derivatives on the aboral side of the arms (Fig.3Cc). These expression patterns were consistent with previous reports of *sfrp1/5* and *fzd5/8* expression in the radial nerve cords of *P. miniata,* and in the ambulacral ectoderm of *P. japonica* for *sfrp1/5* [6,7]. Both genes have additional expression at the tip of the tube feet in *P. miniata*, something we did not observe in *A. squamata* [6].

Next, we investigated the expression profile of the two transcription factors *six3/6* and *nkx2.1*. At the early juvenile stage, *six3/6* showed broad expression in most of the oral side of the disk (Fig.3Da). By the mid-juvenile stage, its expression became restricted to the circumoral nerve ring and the medial part of the ectoneural neuroepithelium in the radial nerve cords (Fig.3Da). Low level of *six3/6* expression was detected in the youngest segments and was more prominently expressed in the brachial ganglia of older segments, where it also extended more laterally from the midline and spanned the entire oral-aboral thickness of the neuroepithelium (Fig.3Dc,Dd). In addition, *six3/6* expression extended laterally at the base of each pair of brachial tube feet, in a region corresponding to the podial ganglia (Fig.3Db). This medial expression pattern of *six3/6* in *A. squamata* was consistent with its expression in regenerating arms of the ophiuroids *A. filiformis* [37], in the radial nerve cords of the asteroid *P. miniata* [6], and in the ambulacral ectoderm of the echinoids *P. japonica* and *Heliocidaris erythrogramma* [7,62] and the crinoid *Anneissia japonica* [63]. Lateral extension of the *six3/6* expression towards the base of the brachial tube feet also appears to be present in *P. miniata*, *H. erythrogramma* and *A. japonica* [6,62,63], indicating a highly conserved expression pattern for *six3/6* across echinoderm classes. The only notable exception seems to be the echinoid *Strongylocentrotus purpuratus*, in which *six3/6* is reported in the tube feet of the echinoid, but not in the radial nerve cords [64].

Finally, *nkx2.1* was expressed in the ectoneural part of the radial nerve cords and in the circumoral nerve ring at all three stages considered. Within the circumoral nerve ring, *nkx2.1* was restricted to discrete regions of the radial and inter-radial ganglia (Fig.3Aa). Similarly, its expression was not continuous along the arm midline and was restricted to discrete regions of each brachial ganglion, but absent from interganglionic regions (Fig.3Ea). In the brachial ganglia, *nkx2.1* exhibited a complex stereotypical expression pattern with the most medial expression at the distal part of the ganglia, while in the proximal region its expression was offset to the sides of the midline (Fig.3Ab). In both cases, the expression of *nkx2.1* spanned the entire oral-aboral thickness of the neuroepithelium (Fig.3Ac,d) and largely overlapped with the distribution of *elav*+ neurons within the ganglia (Fig.3F). Interestingly, this expression pattern contrasted with the other anterior head markers that exhibited continuous expression through the ganglionic and interganglionic regions of the ectoneural neuroepithelium. This was also different from its expression in *P. miniata,* in which it is expressed throughout the entire length of the radial nerve cords and circumoral nerve ring neuroepithelium [6].

**Figure 3:**
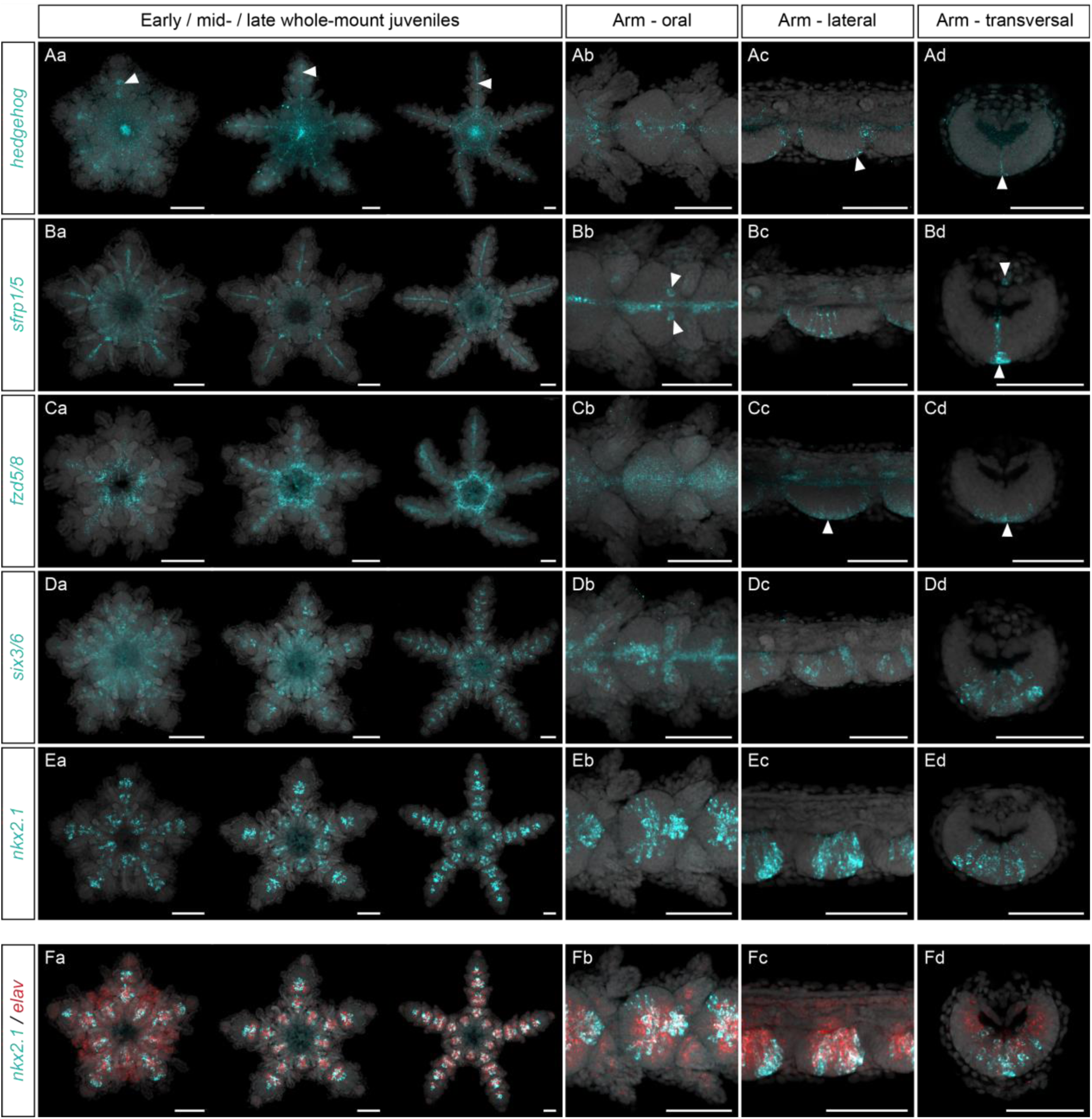
Expression of anterior head markers. Single HCRs for *hedgehog* (**A**), *sfrp1/5*, (**B**), *fzd5/8* (**C**), *six3/6* (**D**), *nkx2.1* (**E**) and double HCRs for *nkx2.1* + *elav* (**F**) in *Amphipholis squamata* whole-mount early (left panel), mid-(middle panel) and late (right panel) juveniles viewed from the oral side (**Aa**, **Ba**, **Ca**, **Da**, **Ea**, **Fa**), and in detailed oral (**Ab**, **Bb**, **Cb**, **Db**, **Eb**, **Fb**), lateral (**Ac**, **Bc**, **Cc**, **Dc**, **Ec**, **Fc**) and transversal (**Ad**, **Bd**, **Cd**, **Dd**, **Ed**, **Fd**) views of brachial segments. Transversal views show sections through the proximal regions of a brachial segment. Arrowheads pinpoint specific expression domains. All samples are counterstained with DAPI (grey) to mark cell nuclei. Scale bars: 100 µm.

### Expression of posterior head markers

Posterior head markers are predominantly expressed in the epidermis covering the tube feet in *P. miniata*, although most of them also overlap with anterior head markers in the medial ambulacral ectoderm, with expression in the radial nerve cords and circumoral nerve ring as well [6]. These genes include the transcription factors *irx*, *dmbx*, *barH*, *otx* and *pax6*, which are all involved in the patterning of the vertebrate forebrain and midbrain and in the hemichordate posterior proboscis and collar [12,57,65,66]. In *A. squamata*, we found that *irx* had the broadest expression domain of all the genes examined in this study, with an expression spanning multiple anatomical structures. This was similar to *P. miniata*, in which *irx* also shows a broad expression in several tissues [6]. In *A. squamata*, *irx* was consistently expressed at the early, mid-and late juveniles in the circumoral nerve ring and the radial nerve cords (Fig.4Aa). Following the early juvenile stage, *irx* expression in the radial nerve cords was markedly stronger in the two most distal arm segments than in the rest of the arm (Fig.4Aa). In the radial nerve cords, *irx* was broadly expressed in the whole ectoneural neuroepithelium, but was absent from the hyponeural neuroepithelium (Fig.4Ab-d). Outside the radial nerve cords, *irx* expression in ectoderm derivatives included the podial ganglia and the epidermis of the brachial tube feet (Fig.4Ab-d). Finally, *irx* also showed additional expression domains in mesoderm derivatives on the aboral side of the arm and in the spines (Fig.4Ac,d), consistent with its expression in the hydrocoel in *P. miniata* [6].

The two next transcription factors considered, *dmbx* and *otx*, shared many similarities in their expression domains. At the early juvenile stage *dmbx* had a diffuse expression in the oral region of the disk (Fig.4Ba). By the mid-juvenile stage, it became restricted to the circumoral nerve ring and the radial nerve cords (Fig.4Ba). Its expression in the radial nerve cords was higher in sub-terminal segments, and less distinguishable in older proximal segments. Within each segment, *dmbx* was predominantly expressed in the interganglionic region of the ectoneural neuroepithelium, but was largely absent from the brachial ganglia (Fig.4Bb-d). In addition, it was expressed in the proximal part of the brachial tube feet epidermis. As reported previously [67], *otx* had a very similar expression pattern to *dmbx*, although it differed in some aspects. First, *otx* started being clearly expressed at the early juvenile stage in the brachial tube feet and persisted all the way through to the late juvenile stage even in proximal segments (Fig.4Ca). It also extended more laterally than *dmbx* in the brachial tube feet epidermis, but not all the way to their extremities (Fig.4Cb-d). In *P. miniata*, both *dmbx* and *otx* have expression domains that span both the radial nerve cords and the tube feet epidermis, but *dmbx* is mostly expressed in the radial nerve cord while *otx* is mostly expressed in the tube feet epidermis. In *A. squamata*, these two genes retained both expression in the radial nerve cords and the brachial tube feet epidermis, but this latter domain was predominant for both of them. Interestingly, *dmbx* and *otx* expression in the radial nerve cords of *A. squamata* was mostly restricted to the interganglionic regions at the junction between the pairs of brachial tube feet, while they were largely (*dmbx*) or completely (*otx*) absent from the brachial ganglia, thus being mutually exclusive with *nkx2.1* expression (Fig.4F). This was a significant difference with *nkx2.1* and *otx* expression in *P. miniata*, in which these two genes overlap [6]. In addition to its expression domain in *P. miniata*, *otx* has been surveyed in developing adult body plan of the largest sample of echinoderm species, and shows a consistent trend of expression on the lateral sides of the radial nerve cords and in the tube feet epidermis, including in the echinoids *P. japonica* [26] and *Holopneustes purpurescens* [68,69]. On the other hand, *otx* expression appears more divergent in the asteroid *Parvulastra exigua* where it seems to be expressed mostly in the radial nerve cords, but has not been reported in the tube feet [70], while in the echinoid *H. erythrogramma* and the crinoid *A. japonica* it is present the epidermis of the tube feet, but was not reported in the radial nerve cords [63,71].

**Figure 4:**
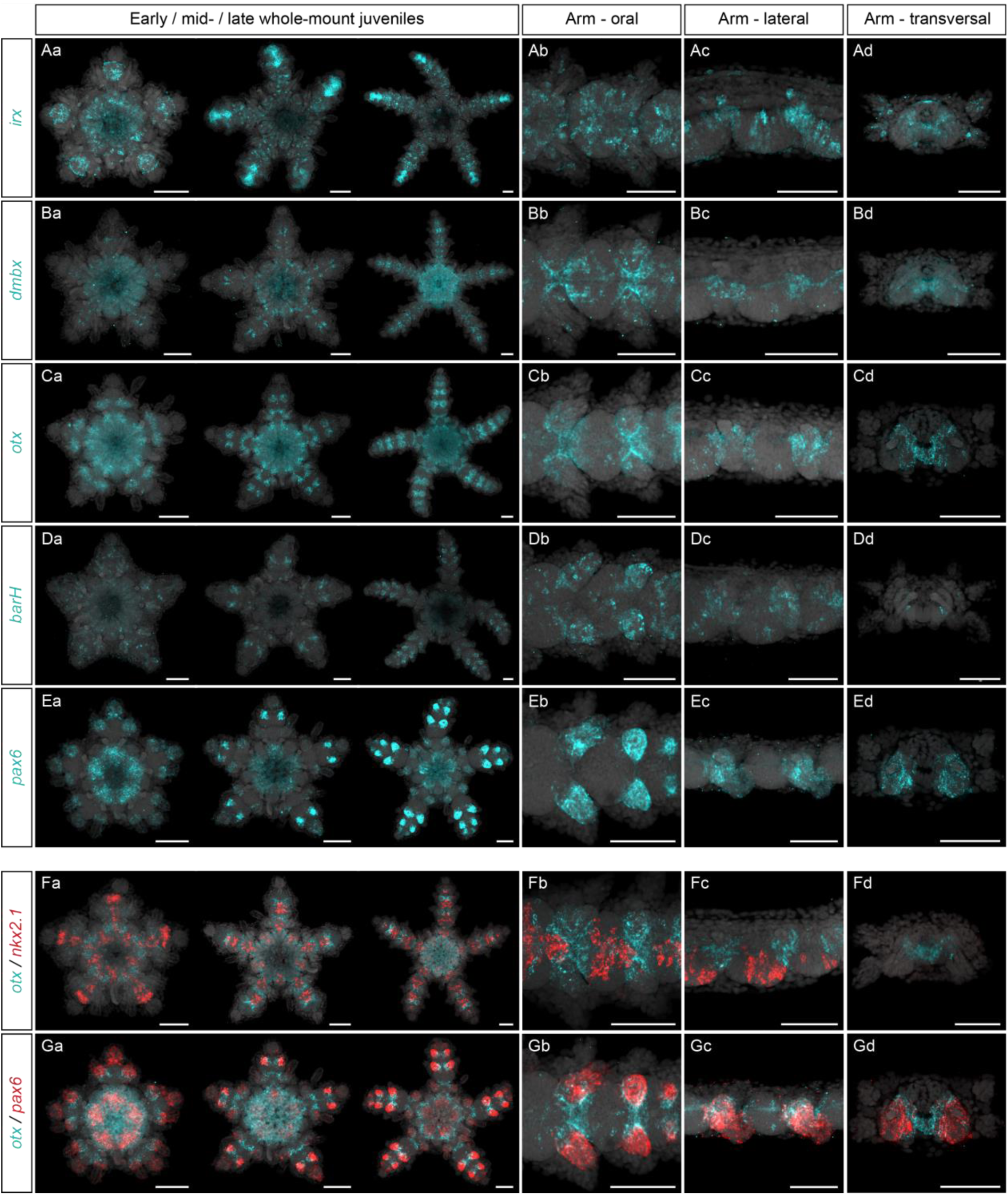
Expression of posterior head markers. Single HCRs for *irx* (**A**), *dmbx*, (**B**), *otx* (**C**), *barH* (**D**), *pax6* (**E**) and double HCRs for *otx* + *nkx2.1* (**F**) and *otx* + *pax6* (**G**) in *Amphipholis squamata* whole-mount early (left panel), mid-(middle panel) and late (right panel) juveniles viewed from the oral side (**Aa**, **Ba**, **Ca**, **Da**, **Ea**, **Fa**, **Ga**), and in detailed oral (**Ab**, **Bb**, **Cb**, **Db**, **Eb**, **Fb**, **Gb**), lateral (**Ac**, **Bc**, **Cc**, **Dc**, **Ec**, **Fc**, **Gc**) and transversal (**Ad**, **Bd**, **Cd**, **Dd**, **Ed**, **Fd**, **Gd**) views of brachial segments. Transversal views show sections through the distal regions of a brachial segment. All samples are counterstained with DAPI (grey) to mark cell nuclei. Note that staining in the gut for *dmbx* and *otx* HCRs results from aspecific background signal. Scale bars: 100 µm.

Finally, *barH* and *pax6* also had largely similar expression domains. *BarH* was first detected at the early juvenile stage in scattered cells of the oral region of the disk (Fig.4Da). Later on, at the mid- and late juvenile stages *barH* expression persisted in scattered cells along the ectoneural part of the radial nerve cords, but also expressed in the epidermis of the brachial tube feet (Fig.4Da-Dd). There, *barH* expression extended all the way up to the tip of the tube feet, unlike *dmbx* and *otx*. Besides *A. squamata*, *barH* expression in echinoderms has only been investigated in *P. miniata*, but shows a very consistent pattern with expression mostly in the tube feet epidermis and in scattered cells in the radial nerve cords [6]. Compared to *barH*, *pax6* was completely absent from the midline of the radial nerve cords and its expression was exclusively restricted to the epidermis of the brachial tube feet, where it was already expressed at the early juvenile stage (Fig.4Ea). At this stage, *pax6* was also expressed in the terminal segment, but this domain did not persist in older stages. Similarly to *barH*, *pax6* expression in the tube feet epidermis extended all the way to the tip, resulting in a much more lateral expression domain than more medial tube feet genes such as *otx* (Fig.4G). This expression domain in *A. squamata* confirms that *pax6* appears restricted to the tube feet epidermis across all echinoderm classes [6,37,62–64,70].

### Expression of head-trunk boundary markers

Head-trunk boundary markers are predominantly expressed at the edges of the ambulacral ectoderm in *P. miniata*, outlining the tube feet epidermis [6]. This category includes the transcription factors *gbx*, *hox1* and *pax2/5/8*, which are known to pattern the midbrain-hindbrain boundary and the collar-trunk boundary in vertebrates and hemichordates, respectively [57,58,72,73]. In *A. squamata*, we found that *gbx* was expressed in the early juvenile stage in the ectoneural part of the developing radial nerve cords (Fig.5Aa,Ab). Specifically, it was expressed in the most lateral areas of the brachial ganglia, and was more strongly expressed in the recent brachial segments, while its expression was reduced in older segments (Fig.5Aa,Ab). In both cases, its expression spanned the entire oral-aboral thickness of the ectoneural neuroepithelium (Fig.5Ac,Ad). Importantly, *gbx* expression in the brachial ganglia was mutually exclusive with the expression of more medial genes like *six3/6* (Fig.5D). Although it was not detected until the mid-juvenile stage (Fig.5Ba), *hox1* expression was similar to *gbx* (Fig.5E) at the most lateral areas of the brachial ganglia, spanning the entire oral-aboral length of the ectoneural neuroepithelium (Fig.5Ba-Bd). Its expression was also mutually exclusive with the expression of medial genes such as of *nkx2.1* (Fig.5F). However, unlike *gbx*, *hox1* was segregated into two clear clusters of cells on each side of the ganglia, one large cluster in the distal part of the ganglia and a much smaller cluster in the proximal part (Fig.5Ba,Bc). The expression of *gbx* and *hox1* at the margins of the brachial ganglia in *A. squamata* appeared very different from its expression in other echinoderm species. In the asteroid *P. miniata*, these two genes are expressed at the margin of the ambulacral ectoderm, but not in the radial nerve cords themselves from which they are separated by the tube feet [6]. In the echinoid *P. japonica*, *gbx* expression has been reported in the mesoderm but not in the ectoderm, and similarly *hox1* expression in the holothuroid *Apostichopus japonicus*, was reported in the digestive tract but not in ectoderm derivatives [27]. On the other hand, *hox1* expression in *P. japonica* clearly outlines the ambulacral ectoderm of the rays II and IV, much more similarly to *A. squamata* and *P. miniata*, but shows a very different expression pattern in the rays I, III and V [26]. However, these differences of gene expression between rays in *P. japonica* must be interpreted in the context of irregular echinoids, in which a secondary bilateral symmetry is superimposed to the pentaradial symmetry [74].

**Figure 5:**
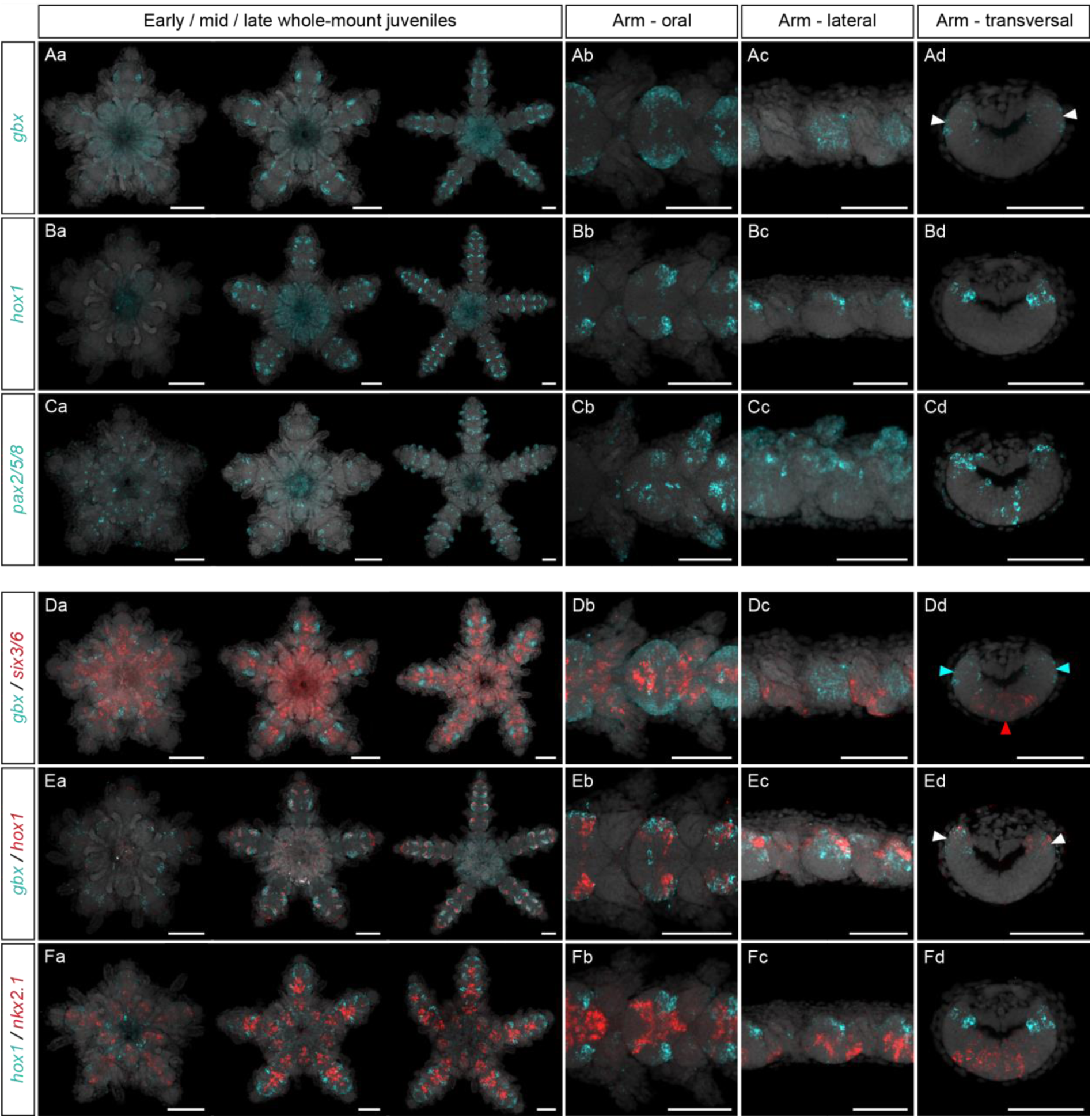
Expression of head-trunk boundary markers. Single HCRs for *gbx* (**A**), *hox1*, (**B**), *pax2/5/8* (**C**) and double HCRs for *gbx* + *six3/6* (**D**), *gbx* + *hox1* (**E**), and *hox1* + *nkx2.1* (**F**) in *Amphipholis squamata* whole-mount early (left panel), mid- (middle panel) and late (right panel) juveniles viewed from the oral side (**Aa**, **Ba**, **Ca**, **Da**, **Ea**, **Fa**), and in detailed oral (**Ab**, **Bb**, **Cb**, **Db**, **Eb**, **Fb**), lateral (**Ac**, **Bc**, **Cc**, **Dc**, **Ec**, **Fc**) and transversal (**Ad**, **Bd**, **Cd**, **Dd**, **Ed**, **Fd**) views of brachial segments. Transversal views show sections through the proximal regions of a brachial segment. Arrowheads pinpoint specific expression domains. All samples are counterstained with DAPI (grey) to mark cell nuclei. Scale bars: 100 µm.

Finally, the expression of *pax2/5/8* is more complex. It began at the early juvenile stage with a punctate pattern in the oral region of the disk (Fig.5Ca). Later, at the mid- and late juvenile stages, distinct expression domains became clear, including in several regions of the radial nerve cords, in the epidermis of the brachial tube feet, and in the epidermis of the developing spines (Fig.5Ca-Cc). In the radial nerve cords, *pax2/5/8* was expressed in scattered cells dispersed in the medial region of the ectoneural epithelium, and in two symmetrical clusters in the lateral parts of the brachial ganglia (Fig.5Ca-Cd). This broad and intricate expression pattern was reminiscent from *P. japonica*, where it is expressed in the epidermis covering the spine rudiments [7], and from *P. miniata*, where it is expressed in scattered cells of the radial nerve cords, at the edge of the ambulacral boundary, and in the interradial epidermis where are located the spines [6].

### Expression of other Hox genes

Hox genes posterior to *hox1* are absent from ectoderm derivatives and are restricted to the endoderm and mesoderm domains in *P. miniata* [6]. In other bilaterian species, Hox genes are posterior markers typically involved in patterning trunk ectoderm territories [75–79]. A recent study in the ophiuroid *A. filiformis* revealed the presence of a full Hox complement, albeit with important syntenic rearrangements [38]. In *A. squamata*, we were not able to detect *hox6*, *hox11/13a* and *hox11/13c* from our Iso-Seq data, and while we were able to identify *hox3* and *hox11/13b*, we could not detect any expression for these two genes by HCRs. Since adult ophiuroids are characterized by a blind gut and lack both intestine and anus, the absence of *hox11/13a* and *hox11/13b* expression in *A. squamata* appeared consistent with the expression domains of these genes in *P. miniata*, in which they are restricted terminally in the intestine [6]. Similarly, *hox11/13b* is expressed in the intestine of the holothuroid *A. japonicus* and the echinoids *S. purpuratus* and *P. japonica* [24,26,27]. In species with abbreviated development such as the crinoid *Metacrinus rotundus* and the echinoids *P. japonica* and *H purpurescens*, *hox11/13b* was surveyed in larval stages lacking a through gut before the formation of the adult digestive tract, making further comparisons impossible [25,26,80]. On the other hand, *hox11/13b* in echinoids, together with *hox11/13a*, has additional expression domains in the coeloms and the interambulacral ectoderm that have no equivalents in *A. squamata* nor *P. miniata* at the stages investigated [24,26].

For the remaining Hox genes (*hox2*, *hox4*, *hox5*, *hox7*, *hox8*, *hox9/10*), we observed highly diverse expression patterns. *Hox2* was not expressed at the early juvenile stage and was later expressed in few scattered cells of the arm muscles and coeloms but was absent from ectoderm derivatives (Fig.6Aa-6Ad). To our knowledge, this is the first report of *hox2* expression in the adult body plan of any echinoderm. However, since *hox2* is only expressed in late juvenile stages in *A. squamata*, its absence from previous surveys in other species might result from sampling biases [6,25,27]. Similarly to *hox2*, the expression of *hox4* was first detected during the development of the arms and was missing at the early juvenile stages. In mid- and late juveniles, *hox4* was expressed on the aboral side of the lateral epidermis of the three most recent sub-terminal arm segments but was absent from the terminal segment itself (Fig.6Ba-Bd). Importantly, *hox4* was the only gene for which we observed extensive expression in the epidermis. This *hox4* expression domain was strikingly different from its expression in the pharynx in *P. miniata* [6] and the hydrocoel in *P. exigua* [81]. *Hox5* was expressed in early juveniles in scattered cells of the developing radial nerve cords (Fig.6Ca). At later stages, its expression in the radial nerve cords became stereotypical within a clearly defined cluster of cells in the lateral parts of each brachial ganglion (Fig.6Ca-Cd). In late juveniles, *hox5* was also expressed in the developing spines (Fig.6Ca-Cc). This *hox5* expression domain in *A. squamata* was a major difference with *P. miniata*, *A. japonicus*, and *M. rotundus*, in which *hox5* is not expressed in ectoderm derivatives and exclusively restricted to coelomic tissues [6,25,27]. Much like *hox5*, *hox7* was expressed in *A. squamata* in lateral clusters of cells within the brachial ganglia, but its expression started later, at the mid-juvenile stage (Fig.6Da-Dd). Similar to *hox5*, it was also expressed at the late juvenile stage in the developing spines (Fig.6Da,Db). Here again, this *hox7* expression domain in *A. squamata* differs from previous reports of *hox7* expression in the developing adult body plan of other echinoderm species. In *P. miniata* and *A. japonicus hox7* expression is reported in the intestine [6,27], while in echinoids and crinoids it is expressed in the somatocoel of the adult rudiment [24–26]. By contrast, we found that *hox8* and *hox9/10* expression patterns were more consistent with that of other echinoderms. In *A. squamata*, *hox8* expression was initially restricted to the ring canal of the water vascular system (Fig.6Ea), a coelomic compartment deriving from the hydrocoel. Later on, *hox8* expression was downregulated in the ring canal, but persisted in five discrete spots corresponding to the position of the radial muscles at the base of each arm (Fig.6Ea). At the late juvenile stage, *hox8* was also expressed in the proximal part of the coelom in the aboral region of the arm segments, on either side of the position occupied by the oral intervertebral muscles (Fig.6Ea-Ed). This was consistent with *hox8* expression in the coelomic compartments of the asteroid *P. miniata* [6], the echinoids *P. japonica* and *S. purpuratus* [24,26], the holothuroid *A. japonicus* [27] and the crinoid *M. rotundus* [25]. Finally, *hox9/10* expression in *A. squamata* was consistent from early to late juvenile stages in the coelomic compartment occupying the aboral region of the disk and the aboral midline of the arms (Fig.6Fa-Fd). This again was consistent with the reported expression of *hox9/10* in the somatocoel of *P. miniata*, *P. japonica*, *S. purpuratus*, *A japonicus* and *M. rotundus* [6,24–27].

**Figure 6:**
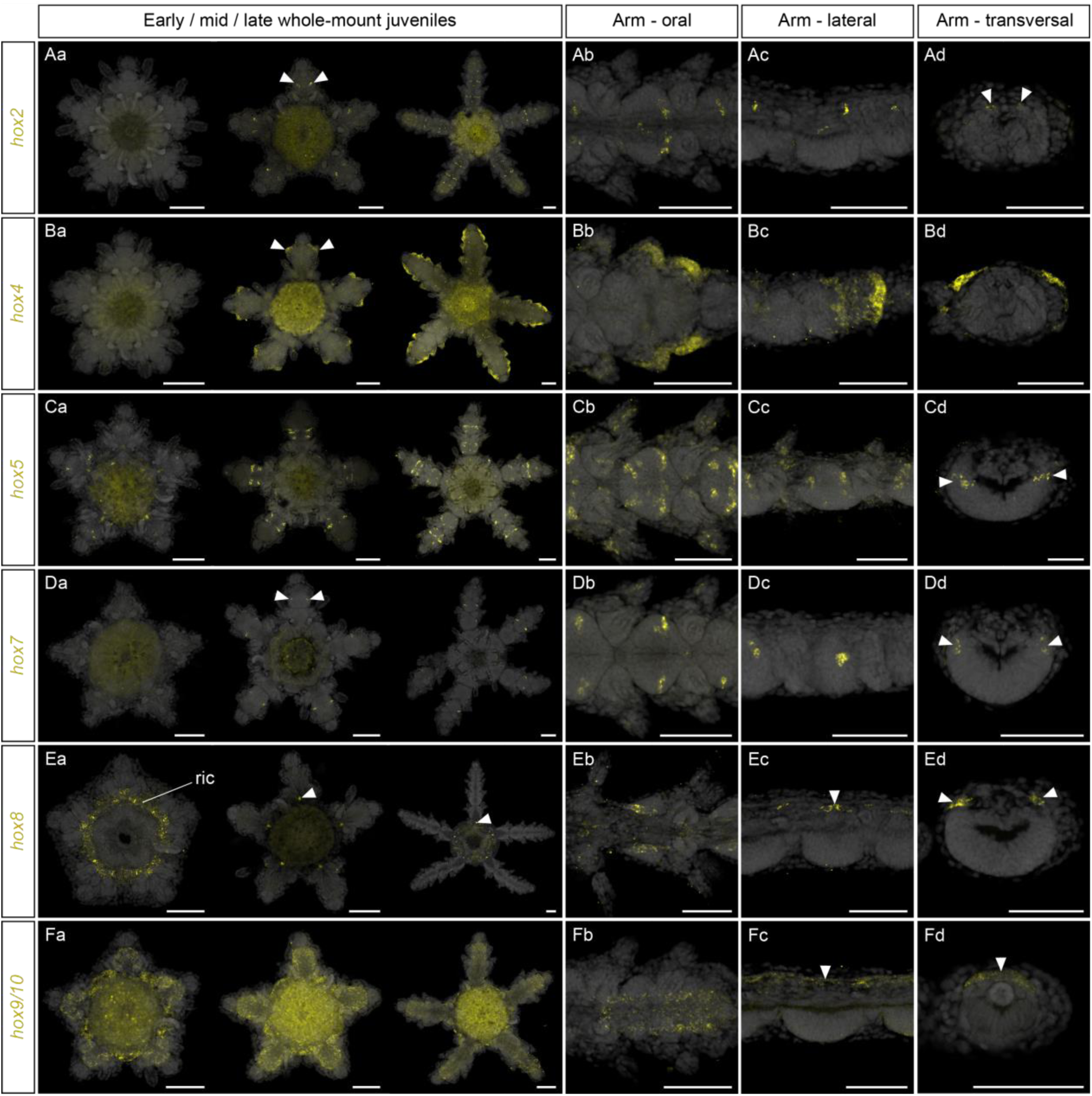
Expression of other Hox genes. HCRs for *hox2* (**A**), *hox4*, (**B**), *hox5* (**C**), *hox7* (**D**), *hox8* (**E**) and *hox9/10* (**F**) in *Amphipholis squamata* whole-mount early (left panel), mid- (middle panel) and late (right panel) juveniles viewed from the oral side (**Aa**, **Ba**, **Ca**, **Da**, **Ea**, **Fa**), and in detailed oral (**Ab**, **Bb**, **Cb**, **Db**, **Eb**, **Fb**), lateral (**Ac**, **Bc**, **Cc**, **Dc**, **Ec**, **Fc**) and transversal (**Ad**, **Bd**, **Cd**, **Dd**, **Ed**, **Fd**) views of brachial segments. Transversal views show sections through the distal (**Ad**, **Bd**, **Fd**) or proximal **(Bd**, **Cd**, **Ed**) regions of a brachial segment. Arrowheads pinpoint specific expression domains. All samples are counterstained with DAPI (grey) to mark cell nuclei. Note that staining in the gut for *hox2*, *hox4* and *hox9/10* HCRs results from aspecific background signal. ric: ring canal. Scale bars: 100 µm.

## Discussion

The relationship between the derived pentaradial symmetry of echinoderms and the bilateral body plan of their bilaterian relatives has puzzled zoologists for over a century [1]. While the identification of conserved axial patterning genes has contributed major insights into understanding metazoan body plan evolution [19,82–86], investigation of echinoderm pentaradial body plans development has only recently begun to provide sufficient comparative data to enable basic axial comparisons with other bilaterians. The limited number of axial patterning studies on adult echinoderms have to date largely focused on the expression of Hox genes in diverse taxa [24–27,69,80,81,87], with the notable exception of ophiuroids. By contrast, the conserved developmental program that patterns anterior territories in bilaterians has only recently been comprehensively investigated in the adult body plan of echinoids and asteroids [6,7], but has been key in developing a new model for relating axial properties of echinoderms to their bilaterian relatives [6]. Here, we extend our understanding of molecular patterning in echinoderm adult body plans by providing a comprehensive summary of AP patterning genes during juvenile development of the ophiuroid *A. squamata*. We found that gene expression data from *A. squamata* are largely congruent with the ambulacral-anterior model previously described in asteroids, but with some significant expression differences that we propose reflect derived ophiuroid morphological adaptations. By analyzing the similarities and differences in gene expression patterns between *A. squamata* and existing datasets from other echinoderm species, we can begin to discriminate between phylum level and class specific regulatory changes involved in adult body plan patterning

Comparing molecular patterning across echinoderm adult body plans has been hampered both by technical challenges and difficulties of comparing data across diverse life history strategies. First, characterizing gene expression in adult rudiments and juveniles is challenging using classical colorimetric *in situ* hybridizations (either whole-mount or on sections) owing to the anatomical complexity of these samples. Fluorescent *in situ* hybridizations, such as HCRs employed here, have largely solved these problems and allow for excellent spatial resolution in anatomically complex samples like echinoderm juveniles. In addition, the heterogeneity of the developmental stages surveyed across species poses another challenge. Investigating post-metamorphic juvenile stages, as it was done here for *A. squamata*, for the asteroid *P. miniata* [6] and for the crinoid *A. japonica* [63] provides a molecular readout of the development of the adult body plan as the ambulacra are extending. Echinoid surveys, however, have historically focused on developmental time points corresponding to the formation of the rudiment that occurs within the larva, prior to metamorphosis. In echinoids, while key ectoderm and mesoderm developmental processes occur during the formation of the rudiment [88–91], other aspects of adult body plan development only take place during and after metamorphosis, such as the formation of the adult digestive tract and the aboral surface of the animal [48,92–94]. Studies during rudiment formation identify regulatory genes involved in the early development of the radial body plan, but the examination of post metamorphic stages is necessary to investigate the full manifestation of the axial properties of the adult. Therefore, cross taxa comparisons need to account for developmental heterochronies that can potentially result in transitory differences in gene expression. Here, we compared the expression patterns observed in *A. squamata* with existing datasets from 10 other species spanning the five extant echinoderm classes (Fig.7A). In the next sections, we discuss how these comparisons refine our understanding of the evolution of echinoderm adult body plan patterning and axial properties.

**Figure 7:**
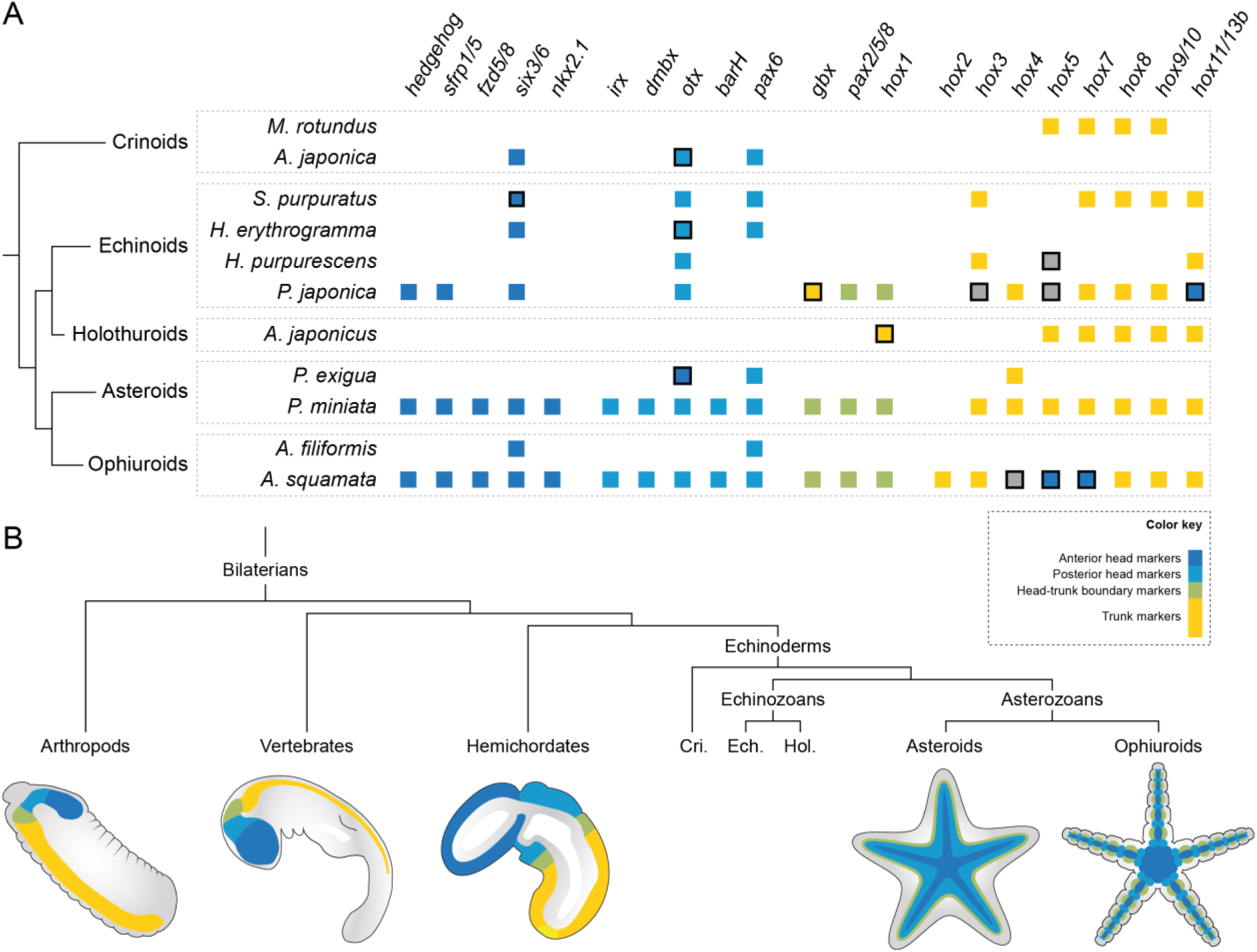
Evolution of axial patterning in echinoderms. **A**, Comparison of the AP patterning genes analyzed in this study in *Amphipholis squamata* juveniles with published datasets from other echinoderm species, including *Metacrinus rotundus* [25], *Anneissia japonica* [63], *Strongylocentrotus purpuratus* [24,64,87], *Heliocidaris erythrogramma* [62,71], *Holopneustes purpurescens* [69,80], *Peronella japonica* [7,26], *Apostichopus japonicus* [27], *Parvulastra exigua* [70,81], *Patiria miniata* [6] and *Amphiura filiformis* [37]. Only genes for which expression was investigated during the development of the adult body plan are reported and expression in embryonic or larval tissues was not considered. Squares indicate genes that have been surveyed, and are colored according to their expression domain (dark blue: medial, light blue: lateral, green: marginal, yellow: internal germ layers, grey: interradial region). Black outlines indicate genes for which the expression pattern does not match the prediction of the anterior-ambulacral model, either for missing important expression domains (e.g. *six3/6* absent from the radial nerve cords in *S. purpuratus*), or being expressed in unexpected territories (e.g. *hox4*, *hox5* and *hox7* expression in ectoderm derivatives in *A. squamata*). Absence of expression for posterior Hox genes is considered as fitting the ambulacral-anterior model. **B**, Schematic representation of the deployment of the AP patterning program in bilateral animals, and in two echinoderm classes following the ambulacral-anterior model. Only expression in ectoderm derivatives is represented. Cri: crinoids, Ech: echinoids, Hol: holothuroids.

### Relationship between molecular patterning and downstream morphology across echinoderm classes

We found that AP patterning genes in *A. squamata* fall into distinct categories largely consistent with previous reports from the asteroid *P. miniata*, and which we referred to as anterior head markers, posterior head markers and head-trunk boundary markers. In *A. squamata*, anterior head markers (*hedgehog*, *sfrp1/5*, *fzd5/8*, *six3/6* and *nkx2.1*) were expressed in the medial region of the radial nerve cords and the circumoral nerve ring. This appears similar to previously described expression patterns in crinoids, echinoids and asteroids [6,7,62,63]. Within the ectoneural part of the radial nerve cords, anterior head markers involved in intercellular signaling pathways (*hedgehog*, *sfrp1/5* and *fz5/8*) and, to a lesser extent the transcription factor *six3/6*, showed continuous expression along the midline of each arm, similar to what was observed in *P. miniata*. By contrast, *nkx2.1* showed discontinuous expression along the proximo-distal axis of the arm, with repeated domains in the ectoneural ganglion of each brachial segment. We suggest that this reflects the upstream role of signaling pathways in setting up the axial properties of the adult body plan, while downstream transcription factors are also involved in the development of particular anatomical structures such as the radial nerve cords. Importantly, radial nerve cords in ophiuroids and echinoids are subepidermal neuroepithelia, while in asteroids and crinoids they are embedded within the epidermis at the bottom of ambulacral grooves [34,35]. This indicates that the conserved expression of the same set of medial genes in the circumoral nerve ring and the radial nerve cords accommodates for significant variability in ontogeny and downstream morphology. Interestingly, the expression of anterior head markers in *A. squamata* neural tissues was restricted to the ectoneural part of the radial nerve cords, but was absent from its hyponeural counterpart. The hyponeural component of the nervous system in echinoderms has been proposed to derive from coelomic mesoderm rather than from ectoderm [55]. Although this idea remains controversial [95,96], the striking discrepancy in the molecular fingerprint of the ectoneural and hyponeural components observed in *A. squamata* suggests marked differences in the development of these two subsystems.

As for anterior head markers, posterior head markers (*irx, dmbx, barH, pax6* and *otx*) also exhibit conserved expression patterns across echinoderm classes, as we observed a strong association of these genes with the development of the tube feet epidermis in *A. squamata* that is consistent with data from other echinoderm taxa [6,7,37,62,63,68,69]. This suggests a highly conserved genetic program for the development of the tube feet in echinoderms – with the only exception of the buccal tube feet that lack *pax6* expression both in *A. squamata* and the echinoid *S. purpuratus* [64].

Our analysis reveals that head-trunk boundary markers exhibit much more variability in their expression domains than anterior and posterior head markers. For example, *gbx* and *hox1* are expressed in different tissues across species. In the asteroid *P. miniata*, *gbx* and *hox1* mark the boundary between ambulacral ectoderm and interradial epidermis, in a territory corresponding to the position of the marginal nerves and the outer limit of the ambulacral grooves [6]. In the echinoid *P. japonica*, *hox1* also outlines the ambulacral ectoderm in the rays II and IV [7]. By contrast, in *A. squamata,* we observed that *gbx* and *hox1* showed similar relative spatial expression to *P. miniata*, laterally compared to anterior and posterior head markers, but within the neuroepithelium of the radial nerve cords and not at the interface between different tissues as in *P. miniata*. Although this represents a significant difference between the two classes, we propose that this may be due to the degree of anatomical divergence in medio-lateral organization between ophiuroids and asteroids, with ophiuroids having no clear equivalent to the ambulacral-interradial boundary of asteroids [1]. This discrepancy suggests flexibility of the patterning system supporting the evolution of class specific anatomies and reflects the role of these genes in providing upstream positional information for the development of the body plan, rather than being tied to the development of particular morphological structures.

Differences in gene expression domains across echinoderm classes are even more marked when considering the Hox genes other than *hox1*. Although *hox8* and *hox9/10* are expressed in coelomic compartments of all echinoderm species investigated – including *A. squamata* – other Hox genes display variable expression domains across classes in either the mesoderm or endoderm. For instance, *hox7* is expressed in the posterior endoderm in both the asteroid *P. miniata* [6] and the holothuroid *A. japonicus* [27], but not in echinoid species [24,26]. However, as mentioned above, time points surveyed in echinoids predate metamorphosis and it is possible that *hox7* turns on later when the intestine of the juvenile starts to develop. Furthermore, Hox gene surveys across several echinoderm species revealed that Hox genes other than *hox1* are largely absent from ectoderm derivatives and are expressed either in endoderm or mesoderm derivatives [6,24–27,69,80,81,87]. Exceptions to this rule were reported in two echinoid species with abbreviated development, *P. japonica* and *H. purpurescens*, where *hox3*, *hox5* and *hox11/13b* have expression domains in the vestibular floor [26,69,80], a territory of the rudiment that has no clear homology within asteroids or ophiuroids. Our results in *A. squamata* add new elements to this list, with the expression of *hox4* in the brachial epidermis and *hox5* and *hox7* in the ectoneural component of the radial nerve cords.

Together, these data indicate a variable degree of coupling between the deployment of upstream patterning genes and downstream morphology across echinoderm classes. Within echinoderms, anterior and posterior head markers show a stronger association with the development of homologous anatomical structures shared across echinoderm classes such as the radial nerve cords and the epidermis of the tube feet. This is consistent with a general association of this molecular program and the formation of neural structures across bilaterians (i.e. the radial nerve cords in echinoderms, the neural plexus of the proboscis in hemichordates, and the forebrain and midbrain in vertebrates), but does not imply anatomical homology between the disparate morphologies regulated by this conserved program across phyla. However, the relationship of head-trunk boundary markers and Hox genes with downstream morphological outputs appears much more variable. The loose coupling between patterning programs, which are responsible for providing axial coordinates during body plan development, and specific morphological outputs appears to be a common theme across metazoans [97–101]. For instance, in hemichordates and vertebrates the same genetic program involving *gbx* controls the formation of the head-trunk boundary in the ectoderm abuting the anterior limit of *hox1* expression, despite the absence of clear anatomical homologies in these regions [58]. Whether these genes have similar functions in establishing anatomical boundaries during the development of the echinoderm adult body plan, such as the ambulacral-interradial boundary in asteroids or the lateral margin of the radial nerve cords in ophiuroids will need to be addressed by future functional studies.

### Evolution of axial properties in echinoderms

Our detailed gene expression map in *A. squamata* allows us to test whether axial patterning in ophiuroids is consistent with the ambulacral-anterior model that proposes a way of comparing the axial properties of adult echinoderms to the anteroposterior axis of bilaterians [6] (Fig.7B). Our findings do not provide evidence supporting the duplication or stacking model. By contrast, despite a few notable differences discussed below, the spatial logic for the deployment of the AP patterning program observed in *A. squamata* is largely consistent with the patterning logic observed in asteroids, from which the ambulacral-anterior model was established. In *A. Squamata*, anterior head markers are expressed in the circumoral nerve ring and along the midline of the radial nerve cords, while head-trunk boundary markers are expressed in the lateral regions of the brachial ganglia. We interpret this as a medio-lateral deployment of the anterior components of the AP patterning program across the radial nerve cords in ophiuroids, which is consistent with the medio-lateral deployment of the same set of genes across the ambulacral ectoderm in the asteroid *P. miniata* [6]. This provides strong evidence to support the ambulacral-anterior model within asterozoans (the clade comprising asteroids and ophiuroids). A broader consideration of anterior patterning gene data from crinoids [63] and echinoids [7,26] also supports this hypothesis. Thus, the reorganization of the ancestral AP patterning program represented by the ambulacral-anterior model likely took place along the stem of the phylum, constituting an ancestral feature of all echinoderm crown-groups (Fig.7B).

There is however a notable difference between the deployment of the AP patterning program described here in *A. squamata* and the ambulacral-anterior model as described in *P. miniata*. In *P. miniata*, posterior head markers (*irx*, *dmbx*, *otx*, *barh*, and *pax6*) are expressed in the epidermis covering the tube feet, and are intercalated between anterior head markers and head-trunk boundary markers that represent more anterior and posterior identities, respectively. Thus, these three territories are organized in the same relative order across the medio-lateral axis of the ambulacral ectoderm as they are along the AP axis in hemichordates and vertebrates. However, in *A. squamata*, the expression of posterior head markers along the arm midline directly abuts the expression of anterior head markers on the side of the brachial ganglia. Posterior head markers, which are expressed in the tube feet epidermis, are offset to the distal part of the brachial segments, instead of being intercalated between anterior head markers and head-trunk boundary markers as in *P. miniata*. This raises the question of which of these two topologies represents the ancestral state for asterozoans. Integrating these molecular data with paleontological evidence allows us to more rigorously test which of the two taxa present the ancestral patterning state of asterozoans. Although reconstructing the evolution of anatomical features in stem asterozoans has proven to be challenging [102,103], there is a consensus that stem ophiuroid body plans were morphologically more similar to extant asteroids than they are to extant ophiuroids [104–106]. Thus, the patterning differences observed in *A. squamata* likely represents a secondary modification in ophiuroids, but not a plesiomorphic trait of asterozoans.

The general absence of Hox expression in the ectoderm of most echinoderm classes led to the idea that these animals lack an equivalent to the trunk of other bilaterian animals [6,7]. However, in *A. squamata* we found that *hox4* is expressed in the arm epidermis while *hox5* and *hox7* are expressed in the ectoneural neuroepithelium. This could indicate that the loss of posterior registry in the ectoderm reached variable degrees across distinct echinoderm classes, with a complete loss in asteroids, but only a partial loss in ophiuroids and possibly echinoids where some Hox genes are expressed in the vestibular floor, as discussed above. Still, we argue that in addition to being less parsimonious, this possibility appears unlikely owing to the absence of collinear Hox expression in the ectoderm of both *A. squamata* and *P. japonica*. We suggest instead that in both cases these expression domains correspond to secondary recruitment of individual Hox genes into novel roles in the ectoderm and do not represent landmarks of a potential trunk territory, corroborating previous interpretations of expression patterns in *P. miniata* and *P. japonica* [6,7]. Importantly, the epidermis in *A. squamata* does not express any of the AP patterning genes investigated in this study, with the exception of *hox4*, indicating that this ectoderm derivative exhibits neither anterior nor posterior registry. This is similar to gene expression data from *P. miniata*, in which the interradial region comprising the epidermis between the ambulacra and on the aboral side of the animal lacks any sign of AP patterning polarity [6]. Similarly, no AP patterning readout was observed in ectodermal derivatives outside the vestibule in the echinoid *P. japonica* [7], although this should be confirmed at later stages by investigating gene expression in the juvenile epidermis. In all three cases, the deployment of the AP patterning program thus appears restricted to the ambulacral ectoderm. In asteroids and crinoids the ambulacral ectoderm, which includes the radial nerve cords, is part of the epidermis at the bottom of opened ambulacral grooves, while in other classes the grooves are covered by skeletal plates and the radial nerve cords are internalized [1,33,34]. The absence of AP registry in the remaining non-ambulacral ectoderm has unclear evolutionary significance, and the origin of this tissue during the development of the juvenile body plan will require further investigations.

While extant echinoderm classes are united by an adult pentaradial body plan, the morphological manifestation of the pentaradial symmetry is highly diverse across classes. Given this profound morphological disparity and the ancient divergence of these classes dating back to the Ordovician [107], patterning divergences are to be expected. With the accumulating wealth of molecular data across extant echinoderm classes and the exquisite fossil record of the phylum, we are now able to readdress the evolution of axial properties both across echinoderm classes and between echinoderms and their bilateral relatives. Future studies will be required to more comprehensively sample across the phylum and validate our prediction that the patterns observed in asteroids and ophiuroids represent stem echinoderm innovations and that the ambulacral anterior model is a valuable tool for exploring echinoderm body plan evolution and diversification. This includes in particular holothuroids and crinoids, for which molecular patterning data have not been extensively investigated. Holothuroids have evolved a secondary bilateral symmetry superimposed to the pentaradial symmetry [108] and have elongated their oral-aboral axis to the point that the homology of their ambulacra with that of other classes is uncertain [109]. Because of these derived anatomical features, important changes in molecular patterning likely occurred in the holothuroid lineage, and it will be key to analyze if the ambulacral-anterior configuration can still be recognized in this class, albeit modified. On the other hand, crinoids constitute the outgroup to other extant classes and are key for any evolutionary scenario as they retain plesiomorphic morphological traits such as the presence of an attachment stalk [110,111]. Thus, comprehensive surveys of molecular patterning in these classes using state of the art gene expression assays are needed.

## Methods

### Animal care

*Amphipholis squamata* adult specimens were collected in Friday Harbor (Washington, USA) and maintained at Hopkins Marine Station (California, USA) on a shallow tank with circulating filtered sea water (FSW) pumped from Monterey Bay (California, USA). The tank was arranged with mud and rocky substrates and overgrown with coralline algae. Every week, it was enriched with a variable amount of *Rhodomonas lens* microalgae culture. *A. squamata* adults reproduced year round on the water table with a peak during spring and summer. For experiments, the largest specimens were retrieved from the water table and used for calcein staining or dissected in a 1:1 mix of filtered sea water and 7.5% MgCl_2_ under a stereoscope. Adult dissections were performed by opening the bursal sacs using a pair of fine tweezers by pinching the proximal part of the epidermis covering the bursae and pulling outward. Developing individuals (embryos, larvae and juveniles) were gently separated from adult tissues, and then processed for *in situ* hybridization. Following dissection, adult specimens were let to recover in a separate tank with circulating FSW for several weeks until they regenerated their bursal sacs, and then put back in the main tank.

### Transcriptome

RNA from adult *A. squamata* arms were isolated using a modified Trizol/RNeasy RNA extraction protocol. In short, samples were homogenized in 1mL of Trizol (Thermofisher) using an extended handle conical tip pestle (Bel-Art Proculture). After vigorously mixing the Trizol homogenate with chloroform, each sample was centrifuged at 10,000g for 18 minutes at 4°C. The aqueous phase was carefully removed and the RNA extract was further purified using a RNeasy Plus Micro Kit (Qiagen). Barcoded PacBio Iso-Seq SMRTbell libraries were constructed using the SMRTbell Express Template Prep Kit 2.0 (PacBio) following the manufacturer’s recommended protocol. The Iso-Seq transcript libraries were bound to the sequencing enzyme using the Sequel II Binding Kit 2.1 and Internal Control Kit 1.0 (PacBio). Sequencing reactions were performed on a PacBio Sequel II System with the Sequel Sequencing Kit 2.0 chemistry. Samples were pre-extended without exposure to illumination for 2 hours to enable the polymerase enzymes to transition into the highly progressive strand-displacing state and sequencing data was collected for 30 hours. Circular consensus sequencing reads were generated from the data using the SMRT Link Version 8.0. For each HiFi read file generated, the data was demultiplexed using lima. Each read file was then refined to include only full length non-chimeric reads. The sequence dataset consisting of all Iso-Seq reads from different samples were combined, clustered and collapsed to reduce gene redundancy while maintaining the highest possible level of gene completeness using CD Hit software tool [112].

### Orthologue identification

Orthologues of genes analyzed by *in situ* hybridizations were identified from the transcriptome by reciprocal best blast hit and validated by phylogenetic trees (Supplementary Material 6). Nucleotide sequences for these transcripts were deposited at GenBank and accession numbers are provided in Supplementary Table 1. Trees were calculated with both the Maximum likelihood using RAxML v.8.2.12 [113] and Bayesian inference using MrBayes v.3.1.262 [114]. For maximum likelihood trees, the robustness of each node was estimated by bootstrap in 1000 pseudoreplicates. For Bayesian inference, trees were calculated in 1,000,000 generations with sampling of trees every 100 generations and a 25% burn-in. *Nkx2.1* sequence variants were identified by blast and a phylogenetic tree was built using EMBL-EBI Clustal alignment tool [115].

### Calcein staining

For calcein stainings, large adult *A. squamata* were collected from the water table and transferred in FSW containing 5 mg.mL^-1^ dissolved calcein, a calcium analogue that is incorporated in the developing skeleton [48]. The animals were incubated for two to three weeks in the dark at 14°C with constant oxygenation, and the calcein-FSW was renewed manually every two days. After two weeks, animals were dissected as described above. Developing individuals which incorporated calcein in their endoskeleton were fixed in FSW containing 3.7% formaldehyde for one hour at room temperature. They were then washed successively in phosphate buffer saline (PBS) containing 0.5% Tween-20 (PBST) and deionized water, and incubated in 50% tetrahydrofuran (Sigma-Aldrich) overnight at 4°C to remove lipids [116]. Samples were then washed successively in deionized water and PBST, and then stained in PBS containing 1:1000 DAPI (Invitrogen) overnight at 4°C before being mounted in a refractive index matching mounting solution (50% weight/volume sucrose, 25% weight/volume urea, 25% weight/volume quadrol) modified from the CUBIC clearing protocol [117].

### Fluorescent *in situ* hybridization

Antisens DNA probes were generated following the probe-split design of HCR v3.0 [118] using HCR 3.0 Probe Maker [119], with adjacent amplification sequences. Probe sets were then ordered as “oligo pools” (Integrated DNA Technology) before being re-suspended in nuclease-free water at a final concentration of 0.5µM. Samples were incubated in fixation buffer (1X phosphate buffered saline (PBS), 0.1M MOPS, 0.5M NaCl, 2mM EGTA, 1mM MgCl2) containing 3.7% formaldehyde overnight at 4°C and then dehydrated in methanol for storage at -20°C. After storage, the samples were rehydrated in deionized water and incubated in 50% tetrahydrofuran (Sigma-Aldrich) overnight at 4°C to remove lipids [116]. Following lipid removal, the samples were washed extensively first in deionized water, and then in PBST, before being permeabilized in detergent solution (1.0% SDS, 0.5% Tween-20, 150 mM NaCl, 1 mM EDTA (pH 8), 50 mM Tris-HCl at pH 7.5) for one hour. Samples were then extensively washed in PBST, and then in 5X saline sodium citrate buffer containing 0.1% Tween-20 (SSCT), before being pre-hybridized in hybridization buffer (Molecular Instruments) for one hour at 37°C. Probes were added to the hybridization buffer at a final concentration of 0.05 µM and the samples were let to hybridize at 37°C overnight under gentle agitation. Following hybridization, samples were washed 4 times 30’ in probe wash buffer (Molecular instruments) at 37°C and then in 5X SSCT at room temperature. They were then pre-amplified in amplification buffer (Molecular Instruments) for 30’. For double HCR and immunohistochemistry, anti FMRF-amide antibody (Immunostar #20091) produced in rabbit was added to the amplification buffer at a final concentration of 1:200. Meanwhile, H1 and H2 components of the HCR amplifiers (Molecular Instruments) were incubated separately at 95°C for 90”, cooled down to room temperature in the dark and then pooled together before being added to the amplification buffer at a final concentration of 60 nM. The amplification reaction was performed overnight. Samples were then extensively washed in 5X SSCT and PBST, and incubated in PBST containing 1:1000 DAPI (Invitrogen) overnight at 4°C. In the case of double HCR and immunohistochemistry, an anti-rabbit secondary antibody coupled to Alexa488 (Sigma-Aldrich) was added at this step at a final concentration of 1:500. Finally, the samples were washed in PBST and transferred to a refractive index matching mounting solution (50% weight/volume sucrose, 25% weight/volume urea, 25% weight/volume quadrol).

### Image acquisition and processing

Images of developing embryos, larvae and juveniles were acquired using a Zeiss Imager A2 equipped with a differential interference contrast setup and a Canon DSLR camera. For large samples, multiple images were tiled together using the automated layer alignment tool in Adobe Photoshop v.12.0.4. Images of calcein stained samples and HCRs were acquired using either a Zeiss LSM700 or a Zeiss LSM900 confocal microscope. Series of optical sections were taken with a z-step interval ranging from 1 to 4 µm depending on sample thickness and signal distribution. Multichannel acquisitions were obtained by sequential imaging. For large samples, multiple images were tiled together using the tiling tool in Zen Blue v.3.8. Optical sections spanning regions of interest were then compiled into maximum intensity z-projections and processed using ImageJ v.1.52g [120].

## Acknowledgements

The authors thank Jeffrey Thompson, Imran Rahman, Maria Byrne, and the members of the Lowe and Rokhsar laboratories for helpful discussions. This work was supported by a Chan Zuckerberg BioHub funding to D.S.R and C.J.L.

## Author contribution

L.F. performed the experiments. P.P. and D.R.R. generated the transcriptome. L.F., D.S.R., and C.J.L. conceived the study, analyzed the results, and wrote the manuscript.

